# Balancing Selectivity and Generality in Object Recognition through Structured Interconnectivity

**DOI:** 10.1101/2024.08.17.608404

**Authors:** Yiyuan Zhang, Jirui Liu, Jia Liu

## Abstract

Balancing selectivity and generality in object recognition is a significant challenge, as it requires the ability to discern fine details that set objects apart while simultaneously embracing the common threads that classify them into one single category. Here we investigated how the brain addresses this challenge by examining the relationship between the interconnectivity of neural networks, the dimensionality of neural space, and the balance of selectivity and generality using neurophysiological data and computational modeling. We found that higher interconnectivity in the TEa of macaques’ IT cortex was associated with lower dimensionality and increased generality, while lower interconnectivity in the TEO correlated with higher dimensionality and enhanced selectivity. To establish the causal link, we developed a brain-inspired computational model formed through Hebbian and anti-Hebbian rules, with wiring length constraints derived from biological brains. The resulting structured interconnectivity created an optimal dimensionality of the neural space, allowing for efficient energy distribution across the representational manifold embedded in the neural space to balance selectivity and generality. Interestingly, this structured interconnectivity placed the network in a critical state that balances adaptability and stability, and fostered a cognitive module with cognitive impenetrability. In summary, our study underscores the importance of structured interconnectivity in achieving a balance between selectivity and generality, providing a unifying view of balancing two extreme demands in object recognition.

## Introduction

Imagine the task of recognizing different breeds of dogs, such as a Dalmatian and a Labrador. Selectivity focuses on the unique features – the distinctive spots and graceful gait of the Dalmatian and the sleek body and friendly eyes of the Labrador, while generality does the opposite, emphasizing common characteristics and overlooking these distinct differences. The challenge in object recognition lies in balancing these processes: differentiating the intricate details that set them apart while simultaneously embracing the common threads that bind them into the single category of ‘dog’ (Grill-Spector & Weiner, 2014; Ullman, 2000; Zoccolan et al., 2007). From a traditional neuronal perspective, selectivity is achieved through neurons that are finely tuned to specific features, while hierarchical networks of neurons integrate this detailed information to generalize and identify commonalities among diverse objects (Felleman & Van Essen, 1991; Riesenhuber & Poggio, 1999, 2002; Rolls & Deco, 2001; Rust & DiCarlo, 2010).

Recent advances in population coding provide a new perspective on how the brain achieves the balance between selectivity and generality. In a high-dimensional neural space constructed by a population of neurons, objects sharing the same features form a neural manifold (Chung & Abbott, 2021; Chung et al., 2018; Mitchell-Heggs et al., 2023). The high dimensionality of this manifold allows for rich and detailed encoding of distinct features, enhancing the ability to distinguish between similar objects through sparse coding and fine-grained representation (Cai et al., 2024; Elmoznino & Bonner, 2024; Mitchell-Heggs et al., 2023). Conversely, the low dimensionality focuses on the most salient and common features shared across different instances of an object category, making the representation robust to variations and thus aiding in quick and efficient categorization (Cohen et al., 2020; Jia et al., 2022). In this framework, we suggest that a single neural system can achieve a balance between selectivity and generality by maintaining an intermediate level of dimensionality. This intermediate dimensionality allows the system to capture sufficient details for distinguishing individual objects while being abstract enough to recognize broader categories. Thus, by avoiding the pitfalls of overly detailed or overly simplified representations, an intermediate dimensionality ensures that the neural coding is both versatile and efficient.

One structural factor that influences the dimensionality of the neural space is the interconnectivity among neurons (Recanatesi et al., 2019; Sporns, 2016). High interconnectivity allows neurons to be densely connected, enabling shared features to be easily combined across different objects. This facilitates the creation of more generalized and abstract representations, ultimately reducing the overall dimensionality of the neural manifold (Barak et al., 2013; Bullmore & Sporns, 2009). Conversely, low interconnectivity leads to neurons that are more specialized and less influenced by other neurons, thereby allowing for a higher-dimensional representation, with each dimension encoding specific and unique features of the object (Felleman & Van Essen, 1991). Thus, to achieve an effective balance between selectivity and generality, we suggest that the neural system might employ an optimal level of interconnectivity.

To test our conjecture that the balance of selectivity and generality can be achieved by maintaining an intermediate level of dimensionality through an optimal level of interconnectivity, we analyzed neurons in the TEO and TEa regions of the inferior temporal (IT) cortex in macaques (Fig. 1A) recorded previously by Bao et al. (2020) when the macaques passively viewed a series of objects from different categories and viewing angles (Fig. 1B). We chose the TEO and TEa as the regions of interest for two main reasons. First, abundant evidence indicates that these regions are central to object recognition, possessing both selectivity and generality (Desimone et al., 1984; DiCarlo et al., 2012; Kobatake & Tanaka, 1994; Tanaka et al., 1991; Wachsmuth et al., 1994). Second, previous anatomical studies of the IT cortex have shown a gradient in dendrite spine density from the posterior to the anterior IT cortex (G. N. Elston et al., 2011; Guy N Elston & Rosa, 1998; Guy N Elston & Rosa, 2000). Specifically, neurons in the TEa possess significantly higher dendritic spine counts (approximately 11,000 per neuron) compared to those in the TEO (about 5,000) (Fig. 1A). Higher dendrite spine density suggests a higher degree of connectivity to other neurons, indicating higher interconnectivity (Guy N Elston, 2007; Sala & Segal, 2014). Therefore, we predicted that the TEa, with its higher interconnectivity, constructs a neural representational space with lower dimensionality, which in turn results in lower selectivity but higher generality, as compared to the TEO with lower interconnectivity.

**Fig. 1.**
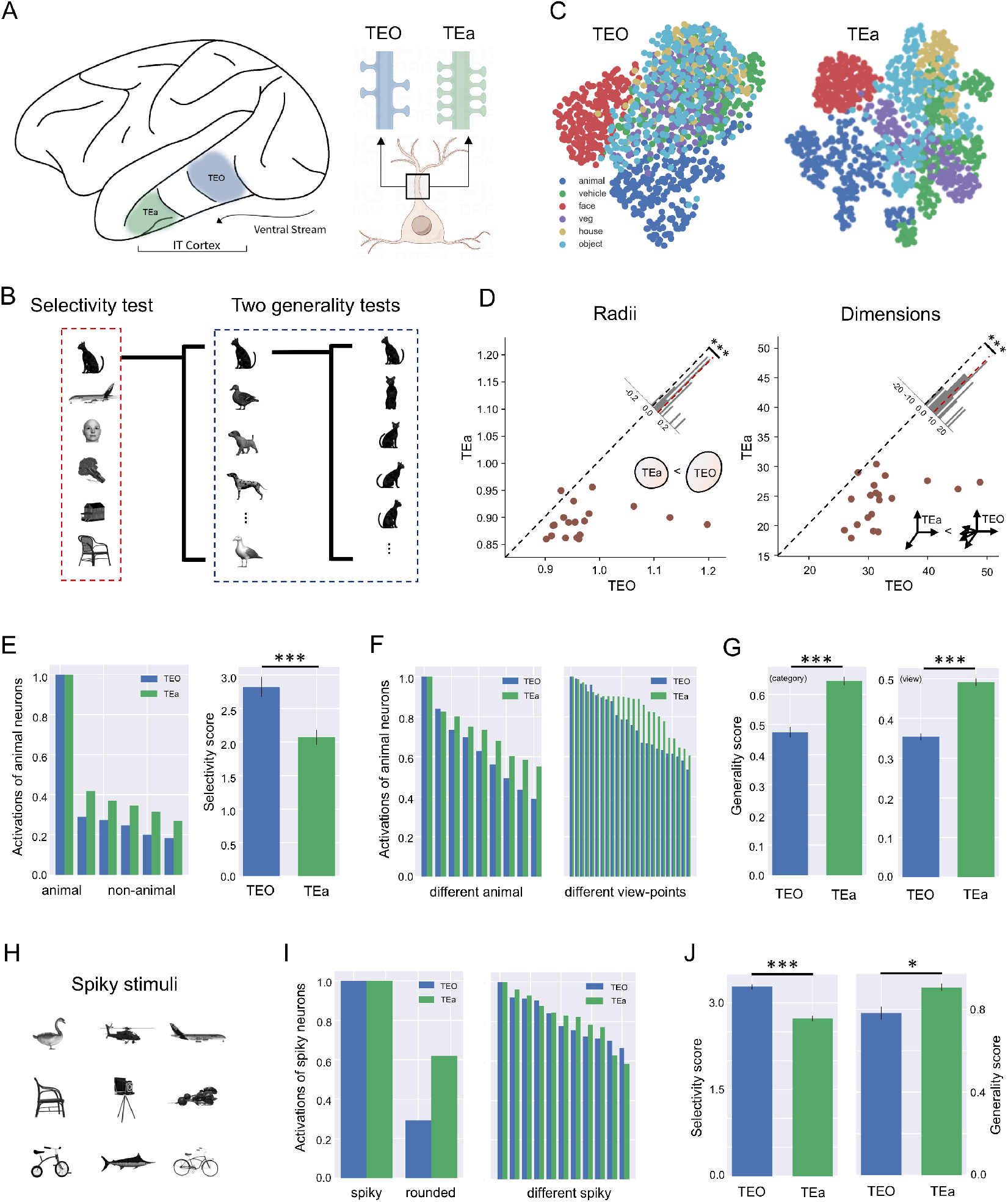
The gradient of selectivity and generality along macaques’ IT cortex. (A) Left: Schematic illustration of the IT cortex, including the TEO and TEa. Right: Schematic illustration of dendritic spines on the dendrites of neurons in the TEO and TEa. Schematic illustrations created with FigDraw (www.figdraw.com). (B) During the recording, macaques passively viewed stimuli from six object categories: animals (10 exemplars), vehicles (11), faces (9), vegetables (7), houses (4), and tools (10). Each exemplar consisted of 24 different views. (C) Neural manifolds of stimuli visualized in a 2-D subspace using UMAP. Each point represents the neural state of one stimulus, with a total of 1224 neural states. Different colors represent different object categories. (D) Radii and dimensions of neural manifolds formed by neurons in the TEO and TEa. Diagonal lines indicate equal radii (left) and dimensions (right). Points below the diagonal lines indicate higher dimensions and larger radii of the TEO compared to the TEa. Each point is the result of a permutation test, randomly sampling 100 neurons from the neuron pools in TEO and TEa, respectively. Histograms in the upper right show the distribution of radii and dimensions. (E) Selectivity. Response magnitudes of animal-responsive neurons in the TEO and TEa to six object categories (left) and corresponding selectivity scores (right). (F) Generality. Response magnitudes of animal-responsive neurons in the TEO and TEa to various animal exemplars (left) and different views of the same exemplar (right). (G) Generality scores of the TEO and TEa in generalizing across different animal exemplars (left) and different views (right). (H) Exemplars of spiky stimuli. (I) Response magnitudes of spiky-responsive neurons in the TEO and TEa to spiky versus non-spiky stimuli (selectivity, left) and to different types of spiky stimuli (generality, right). (J) Selectivity and generality scores of the TEO and TEa in differentiating spiky versus non-spiky stimuli (left) and in generalizing across different spiky stimuli (right). ***: *p* < 0.001; *: *p* < 0.05.

## 2. Results

### 2.1 Gradients of interconnectivity, invariance, and dimensionality in IT cortex

A total of 307 neurons were recorded from the TEO (106 neurons) and TEa (201 neurons) in response to six object categories: animals, vehicles, faces, vegetables, houses, and tools. Each category contains 4 to 11 exemplars, each viewed from 24 different views, which made a total of 1224 stimuli (Fig. 1B). To visualize neural manifolds of these object categories, formed by TEO and TEa neurons, we utilized the Uniform Manifold Approximation and Projection (UMAP) method (McInnes et al., 2018) to project the high-dimensional neural states of all object exemplars into a 2-D subspace (Fig. 1C). Visual inspection of the UMAP projection revealed that in the TEa, neural states of object exemplars were more tightly clustered within each point cloud and more dispersed between them compared to those in the TEO. To quantify this observation, we utilized a method developed by Chung et al. (2018) to calculate the effective dimensions of the manifolds and the radii of point clouds formed by neurons in the TEO and TEa, respectively. Consistent with the visual inspection, the radius of each point cloud in the TEa was significantly smaller than that in the TEO (*p* < .001) (Fig. 1D, left). Critically, the effective dimensionality of the neural manifold in the TEa was significantly lower than that in the TEO (24.19 versus 32.40; *t*(38) = 5.31, *p* < .001) (Fig. 1D, right). This suggests that the TEa, with its higher interconnectivity, constructs a neural representational space with lower dimensionality, embedding the neural states of object categories in more clustered point clouds with smaller radii. Taken together, we observed a gradient of decreased dimensionality along the IT cortex, accompanied by increased interconnectivity.

To explore the gradient of invariance along the IT cortex, we examined the selectivity and generality of these two regions. Among all neurons recorded, 16.9% showed selective responses to a single object category, where object-responsiveness was defined as the average response magnitude to one object category being at least twice the maximum response magnitude to the rest categories. Notably, most of these neurons (78.8%) were specifically tuned to the animal category (Table S1). Accordingly, we focused on the selectivity and generality of the TEO and TEa neurons in response to the animal category. For visual comparison, we ranked the response magnitudes of animal-responsive neurons to animals and non-animal objects in descending order, normalizing the highest responses (i.e., to animals) to 1 (Fig. 1E, left). We found that animal-responsive neurons in the TEO exhibited relatively lower responses to non-animal objects compared to those in the TEa (Fig. 1E, left), indicating a higher level of selectivity in the TEO (*t*(38) = 4.46, *p* < .001) (Fig. 1E, right). Note that the level of selectivity was indexed by the ratio of the average response magnitude to animals to the maximum response magnitude to non-animal objects (see Methods).

A similar analysis was conducted to examine the generality by classifying ten animal exemplars, each consisting of 24 different views, into six basic-level animal categories (i.e., 2 dogs, 1 cat, 1 horse, 3 birds, 1 duck, and 1 fish) (Fig. 1B). Fig. 1F (Left) shows the magnitudes of animal-responsive neurons to different animal exemplars ranked in descending order, with the highest response normalized to 1. We found that neurons in the TEa exhibited higher responses to most animal exemplars compared to those in the TEO, indicating a significantly higher degree of generality in the TEa (*t*(38) = -5.79, *p* < .001) (Fig. 1G, left). Note that the level of generality was indexed by the ratio of the minimum to maximum magnitudes of the animal-responsive neurons to different animal exemplars (see Methods). A similar pattern was found for the generality across different views within exemplars (Fig. 1F, right), indicating that neurons in the TEa showed a significantly higher level of generality across views compared to those in the TEO (*t*(38) = -11.18, *p* < .001) (Fig. 1G, right). Note that the level of generality was indexed by the ratio of the minimum to maximum magnitudes of the animal-responsive neurons to different views (see Methods).

Another index to measure generality is standard deviation, with a lower level of standard deviation among neural responses to animal exemplars indicating a higher level of generality. We found similar results using this index (Fig. S1). Thus, along the gradient of increased interconnectivity, we observed a gradient of increased invariance from selectivity to generality along the IT cortex. Additionally, we examined the selectivity and generality in the TEp, whose neurons exhibit a comparable number of dendritic spines to those in the TEO (i.e., a similar level of interconnectivity) but receive inputs more similar to those of the TEa (G. N. Elston et al., 2011; Guy N Elston & Rosa, 1998; Guy N Elston & Rosa, 2000; Webster et al., 1994). We found that the pattern of dimensions, selectivity and generality in the TEp was more similar to that of the TEO (Fig. S2), supporting the conjecture that interconnectivity plays a critical role in influencing selectivity and generality in object recognition.

The initial analysis focused on categorizing objects into natural categories, such as animate versus inanimate. Here we further examined whether the gradient of increased invariance is also applicable to specific object features across different object categories. Specifically, we re-labelled objects based on the presence of protrusions, categorizing them as either “spiky” (objects with protrusions, Fig. 1H) or “non-spiky” (Bao et al., 2020). In the TEO and TEa, 46.58% of neurons showed a higher response to spiky objects, whereas only 4.23% showed the opposite response pattern. Accordingly, we selected spiky-responsive neurons for further analysis. We found that the spiky-responsive neurons in the TEO exhibited a higher level of selectivity (*t*(38) = 6.25, *p* < .001), whereas the TEa showed a higher level of generality (*t*(38) = -2.56, *p* < .05) (Fig. 1I & J). Thus, the differential preference for selectivity and generality observed in the TEO and TEa extends to specific object features, not just natural object categories, indicating that the gradient of increased invariance along the ventral pathway is a general property of the IT cortex, applicable across various object recognition contexts. In addition, we conducted this analysis for each monkey separately and replicated the main findings in both monkeys, showing that the TEO exhibited significantly higher dimensions, larger radii, higher selectivity, and lower generality compared to TEa (Fig. S3 and S4).

In summary, neurons in the TEa, which possess a higher level of interconnectivity and thus lower dimensionality, compromised their selectivity to their preferred object category to enhance their proficiency in generalizing across different exemplars of the same category compared to those in the TEO. This relationship between interconnectivity and object recognition supports our conjecture that interconnectivity modulates the dimensionality of neural space, which subsequently influences selectivity and generality. However, there are three unresolved issues. First, the relationship between the density of dendrite spines and the degree of interconnectivity is indirect; second, the link between interconnectivity and dimensionality is only associative but not causal; finally, the mechanism by which dimensionality modulates the balance between selectivity and generality remains unclear. To address these issues, we next constructed a brain-inspired neural network aimed at establishing a causal link among interconnectivity, dimensionality, and invariance in object recognition, and elucidating underlying mechanisms.

### 2.2 Brain-inspired neural model to link interconnectivity to functionality

To explore the causal link between interconnectivity and object recognition, we constructed a brain-inspired neural network that models the human ventral temporal cortex (VTC), homologous to macaques’ IT cortex, due to extensive examination on its role in selectivity and generality in object recognition (Downing et al., 2006; Grill-Spector & Weiner, 2014; Tong et al., 2000). This computational model allows for direct manipulating network’s interconnectivity to examine the resultant effects on object recognition.

The construction of this neural network involved four major steps (Fig. 2A). First, we implemented category-responsive regions in a 2-D lattice of a self-organized map (SOM) to simulate human VTC (Aflalo & Graziano, 2006; Cowell & Cottrell, 2013; Doshi & Konkle, 2022; Durbin & Mitchison, 1990; Konkle, 2021; Obermayer et al., 1992; Zhang et al., 2024) (Fig. 2A, left). To do this, we acquired a high-dimensional object space from AlexNet, a pre-trained deep convolutional neural network specifically designed for object recognition (Krizhevsky et al., 2012), and then mapped it onto the 2-D lattice (see Methods). We then fed the stimuli from four categories (face, tools, body, and place) used in the fMRI study of the HCP dataset (Van Essen et al., 2013) into this network to generate four distinct clusters, with neurons in each cluster tuned to one of the object categories (Fig. 2A, right). Previous studies have shown that this 2-D lattice of the SOM exhibits a topological hierarchy and functional specialization similar to those observed in the VTC (Zhang et al., 2024).

**Fig. 2.**
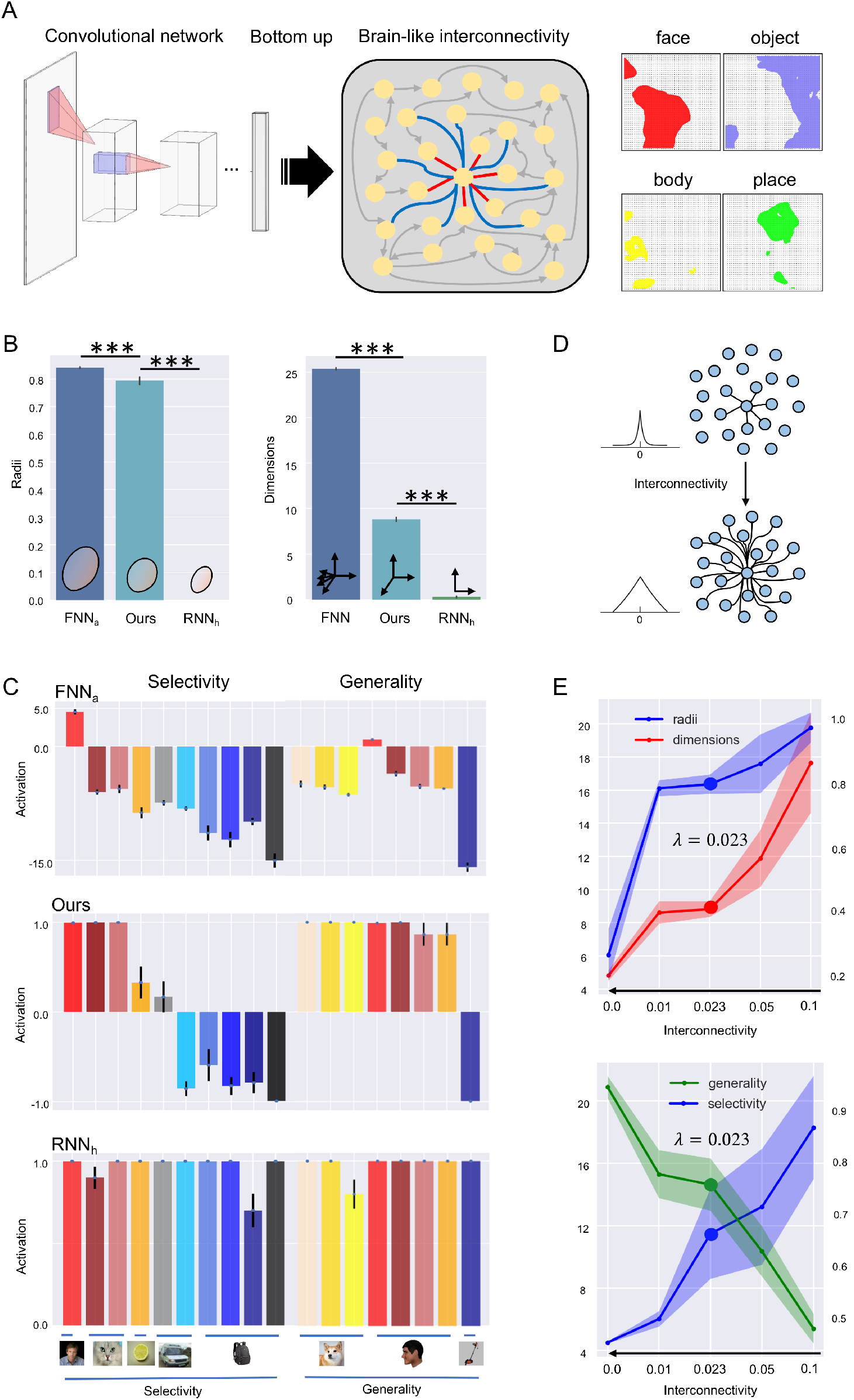
The brain-like computational model. (A) Left: The architecture of the model comprises two main parts: a DCNN encoder, which translates images into vector representations (i.e., object space), and a 2-D lattice of SOM with physical lateral connections to simulate human VTC. Right: The SOM lattice was organized into category-responsive clusters for faces, daily objects, bodies, and places, mirroring the functional specialization observed in the VTC. (B) Radii and dimensions of representational manifolds in three types of neural networks (NN): feedforward NN (FNN_a_), our model, and recurrent NN (RNN_h_). (C) Selectivity and generality in object recognition. Stimuli used for testing selectivity included human faces, faces from different species (dog and cat), objects with shared shapes (lemon), objects with shared configurations (ambulance and airplane), and familiar objects (store, backpack, pitcher, speaker). Stimuli for testing generality included faces from different species (cat, dog, and tiger), human faces from different views (front, profile, cheek, back), and tools as a baseline. Detailed exemplars of these stimuli are provided in Fig. S5. Bar charts show the average activations of the networks with standard deviation. Note that the average activation of our model was derived from neurons in the face cluster after the model had stabilized. (D) Schematic illustration of the parameter *λ* in the exponential distance rule (EDR), a constant that determines the range of lateral connections and thus modulates the interconnectivity of the network. The smaller the parameter λ, the wider the connections between neurons. (E) Top: Radii (blue) and dimensions (red) varied with different levels of interconnectivity in our model. Left y-axis: dimensions; right y-axis: radius sizes (a.u.). Bottom: Selectivity (blue) and generality (green) scores across different levels of interconnectivity. Left y-axis: selectivity scores; right y-axis: generality scores; x-axis: wiring lengths *λ* in a descending order. Shaded areas denote standard deviations. ***: *p* < 0.001.

Second, we implemented the spatial component of network’s interconnectivity by adding physical lateral connections among neurons in the lattice, which are typically absent in standard SOMs. These connections were based on the principle of wiring cost minimization that neuronal interconnections are predominantly local with some long-range connections (Bullmore & Sporns, 2012; y Cajal, 1995), encapsulated by the exponential distance rule (EDR) (Ercsey-Ravasz et al., 2013; Horvát et al., 2016; Theodoni et al., 2022):

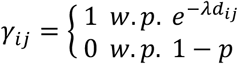

where *γ*_*ij*_ denotes the physical connection between neuron *i* and *j*, and *d*_*ij*_ is the Euclidean distance between them in the lattice. The probability of connection decreases as *d*_*ij*_ increases. In the EDR, the most critical parameter is *λ*, a constant to determines the range of lateral connections. This parameter controls how rapidly the probability of a connection decreases with distance, effectively limiting the influence of neurons on each other based on their spatial separation within the network. In the human temporal lobe, *λ* is estimated to be approximately 0.1 (Theodoni et al., 2022). To calculate the corresponding *λ* in the lattice, we utilized the Symmetric Normalization algorithm (Avants et al., 2008) to warp the lattice of the SOM to the flattened human VTC in the Multi-Modal Parcellation template (Glasser et al., 2016), resulting in a model-corrected *λ* of 0.023 (see Methods). Therefore, *λ* in our model reflects the biological constraint of wiring length in human VTC.

Third, we specified the functional component of interconnectivity using both Hebbian and anti-Hebbian rules (Amit & Amit, 1989; Hopfield, 1982, 1984). The Hebbian rule strengthens excitatory connections between functionally similar neurons, whereas the anti-Hebbian rule establishes inhibitory connections between functionally different neurons:

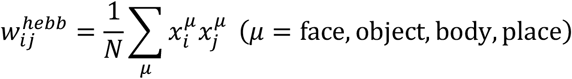

where 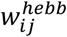 represents the weight between neuron *i* and *j*, determined by their activation states within the *µ*-th object-responsive cluster, and *N* is the total number of neurons. In this formula, 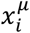 indicates the activity state of the *i*-th neuron for the *µ*-th cluster. For example, if neuron *i* is in the face cluster, then 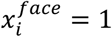, while 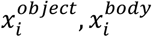, and 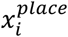 are all set to −1. This configuration creates excitatory connections within the same cluster and inhibitory connections across different clusters. Accordingly, the connection weight *w*_*ij*_ between neuron *i* and *j* is achieved by incorporating both spatial and functional components: 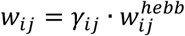, which together determines the structure of network’s interconnectivity.

Fourth, we incorporated temporal dynamics into the network to ensure continuous neural state updates (Fig. S6 A), reflecting the stochastic nature of neural processing (Amit & Amit, 1989; Hertz, 2018). The probability of activation of neuron *i* given condition *b*_*i*_ is a sigmoid function weighted by 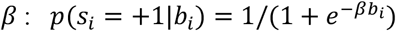. The magnitude of *β* represents intrinsic noise in dynamics, which was set to 100 to simulate a very low-level noise.

In summary, by integrating physical lateral connections governed by EDR (*λ*), Hebbian and anti-Hebbian rules based on functional similarity, and temporal dynamics, we constructed a brain-like structured interconnectivity within the network. By systematically changing the parameter *λ* of wiring length, we can manipulate the network’s level of interconnectivity. This manipulation enables us to examine how changes in the level of interconnectivity affect the dimensionality of the neural space, which in turn influences the balance between selectivity and generality in object recognition.

To evaluate the performance of our model, we compared it with a standard feedforward neural network (e.g., AlexNet of DCNNs, FNN_a_), which lacks lateral connections among neurons, and a standard recurrent neural network (Hopfield Neural Network of RNNs, RNN_h_), characterized by fully interconnected neurons (see Methods). Both feedforward and recurrent neural networks have been widely deployed in object recognition tasks (LeCun et al., 2015; Schmidhuber, 2015). As expected, the FNN_a_ exhibited the highest dimensionality and the largest radius of point clouds (Fig. 2B), indicating a more detailed and complex representation of object features. In contrast, the RNN_h_ displayed the lowest dimensionality and the smallest radius (Fig. 2B), reflecting a more compact and abstract representation. Our model showed intermediate values in both dimensionality and radius, suggesting a balanced representation of object features.

We then evaluated the networks’ ability to recognize faces, a category extensively studied in terms of selectivity versus generality, offering abundant empirical data for comparison. Three types of representative stimuli were chosen for this evaluation. The first type included faces from different species (e.g., dogs’ and cats’ faces) and various view angles (e.g., profile, cheek) to assess the networks’ generality. The second type included non-face objects that share shapes (e.g., lemons) or configurations (e.g., ambulances and airplanes), and the third type included daily objects as familiar as faces (e.g., tools, backpacks), both aimed at evaluating the networks’ selectivity. The FNN_a_ showed extreme selectivity to the human faces it was trained on, with minimal response to other face types or non-face objects (Fig. 2C, top). In contrast, the RNN_h_ showed extreme generality, responding broadly to various faces and non-face objects (Fig. 2C, bottom). Our model, which incorporates structured interconnectivity, struck an ideal balance between selectivity and generality. Specifically, the face cluster in our model responded robustly to all tested face types (i.e., generality) while showing limited response to non-face objects (Fig. 2C, middle). This balanced selectivity and generality mirrors findings from functional neuroimaging studies on the fusiform face area (FFA) in humans, named for its role in face perception (Downing et al., 2006; Tong et al., 2000).

Critically, to investigate the influence of the level of interconnectivity on face recognition, we systematically manipulated the parameter *λ*, which reflects wiring length between neurons (Fig. 2D). Specifically, we varied it to 0.0, 0.01, 0.05, and 0.1, either above or below the value typical of the human cortex (*λ* = 0.023, model-corrected). We found a monotonic increase in both dimensionality and radii as interconnectivity decreased (dimensionality: one-way ANOVA, *F*(4,145) = 168.99, *p* < .001; radii: one-way ANOVA, *F*(4,145) = 382.92, *p* < .001) (Fig. 2E, top). These findings are consistent with observations from macaques’ IT cortex, where higher interconnectivity in the TEa resulted in a lower effective dimensionality and radius, suggesting that interconnectivity plays a causal role in modulating the geometric characteristics of neural manifolds.

In addition, we assessed our model’s selectivity and generality in face recognition by varying solely the parameter *λ* (i.e., different levels of interconnectivity). As expected, the model’s selectivity monotonically increased as interconnectivity decreased, whereas generality showed the opposite trend, decreasing with lower interconnectivity (Fig. 2E, bottom). Critically, the model with a wiring length corresponding to the human cortex (i.e., λ = 0.023) achieved an optimal balance between selectivity and generality. This suggests that our brain is equipped with a neural architecture precisely calibrated for balancing these competing demands, necessitating a dedicated equilibrium between selectivity and generality for daily life object recognition tasks.

### 2.3 Energy distribution across representation manifold

To investigate why with this biologically constraint wiring length, our model achieved the optimal balance between selectivity and generality, we examined the energy distribution across the neural manifold as neural states evolved toward directions of lower energy during the stabilization of the network (Amit & Amit, 1989; Hertz, 2018). To do this, we developed a dimension reduction method, namely parametric Uniform Manifold Approximation and Projection (p-UMAP, see Methods), which parametrizes the non-linear mapping from a high-dimensional space (Fig. 3A, left) to a 2-D representational space (Fig. 3A, middle) using machine learning. Fig. 3A (middle) shows the 2-D representational space after the p-UMAP, where various types of faces were clustered together (i.e., generality) and separated from non-face objects (i.e., selectivity). Then, we calculated the network’s energy for each neural state to visualize the dynamics of the network. This energy is defined as:

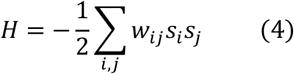

where *s*_*i*_ is the state of neuron *i*, and *w*_*ij*_ is the connection weight from neuron *j* to neuron *i*. Each network state corresponds to an energy value. These energy values were incorporated into the 2-D representational space as a third axis, resulting in a 3-D energy-representational manifold (Fig. 3A, right) (see Methods). This manifold provides a comprehensive view of how neural states evolved, highlighting regions of stability (low energy) and instability (high energy), thereby offering insights into the network’s functional organization.

**Fig. 3.**
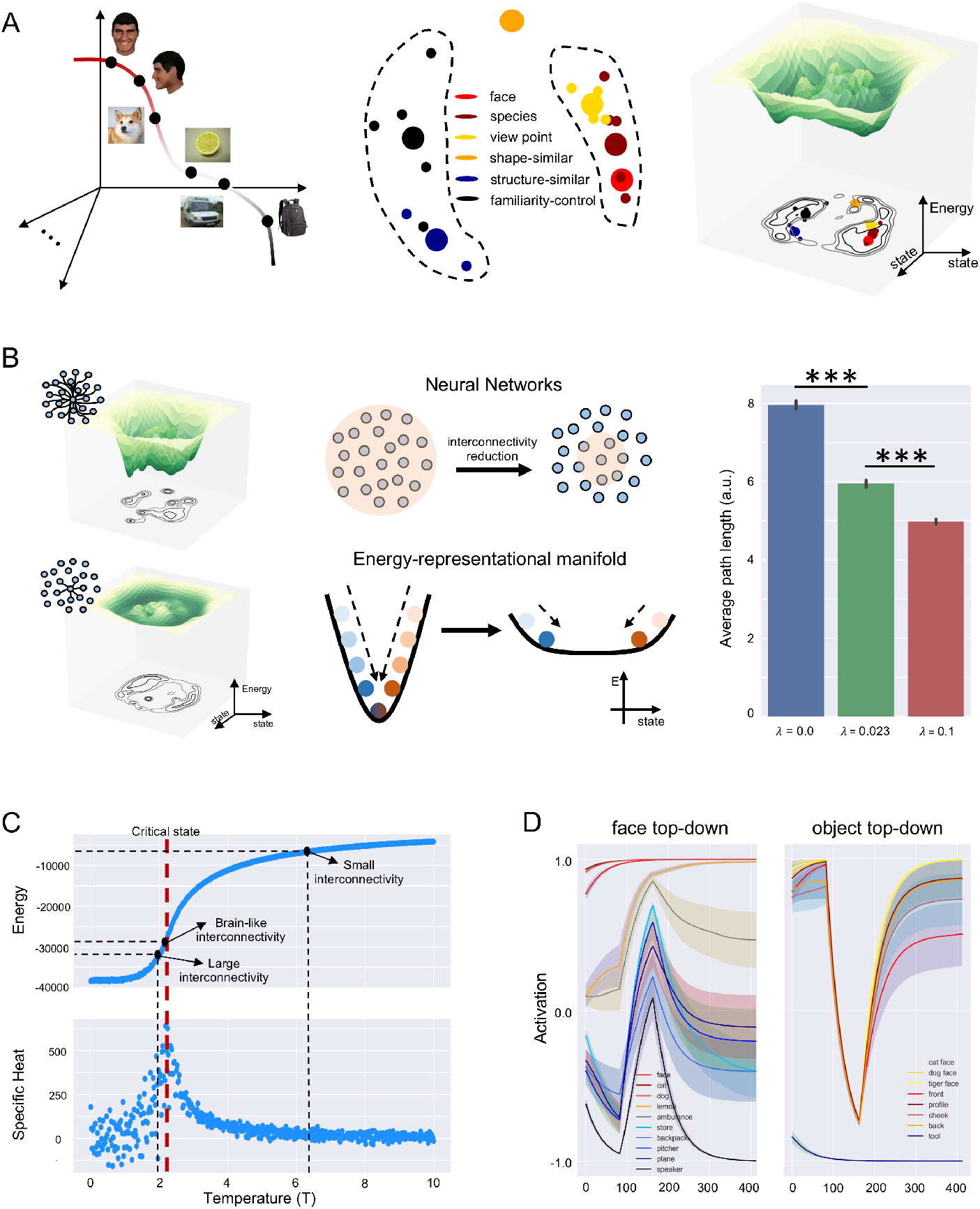
Energy-representational manifold and its characteristics. (A) Left: Schematic illustration of the object manifold in high-dimensional neural space. Face stimuli are located on the red line segment, while tool stimuli are positioned on the black line segment. Objects that share shapes or configurations with faces are situated in between. Middle: The 2-D representational manifold of object categories projected from the high-dimensional neural space via the parametric-UMAP method. Each small dot represents one object category at the basic level (e.g., cat face), and colors of the dots indicate object categories at the conceptual level (e.g., faces of different species). Big dots denote the center of object categories at the conceptual level. Right: Energy of neural states incorporated into the 2-D representational manifold as a third axis, thus forming a 3-D energy-representation manifold. Contour lines projected onto the representational manifold denote energy values, illustrating the shape and elevation of the energy. Contour lines close together indicate a steep slope, while lines spaced further apart indicate a gentler slope. (B) Left: Energy-representation manifolds constructed by models with a longer (λ = 0.0, left) and shorter (λ = 0.1, right) wiring length, respectively. Note that contour lines with a longer wiring length (top) are closer together than those with a shorter wring length (bottom), indicating a steeper slope in the changes of energy in the manifold. Middle: Schematic illustration of the attractor regions in the energy-representation manifolds with different slopes. The steeper the slope, the longer the path that takes the initial neural state to travel to the stable state. Right: Average path lengths of attractor regions in models with different wiring lengths. Our model with the biologically constraint wiring length formed attractor regions that were neither too deep nor too shallow, suggesting a balanced dynamic. (C) Criticality. Hamiltonian value (top) and specific heat (bottom) at different temperatures of the Ising model. The Hamiltonian value for the model with λ = 0.023 lay close to the critical point of the Ising model, whereas models with either a longer or a shorter wiring length resided in either order or chaos regions. (D) Cognitive impenetrability. Left: A top-down signal of faces was applied to all units (indicative of the presence of faces), while inputs were continuously presented objects (e.g., lemon, ambulance, and backpack). Right: A top-down signal of objects was applied to all units (indicative of the presence of objects), while input stimuli were continuously presented various faces. Lines denote time courses of activation averaged across all units in the face cluster along iteration steps, with shaded areas denoting standard deviation. Time courses show the dominant effect of top-down signals; however, once the top-down signals were offset, the time courses resumed to their original trajectories, suggesting the cognitive impenetrability of the model. ***: *p* < 0.001.

For comparison, we also generated energy-representation manifolds for networks with either a longer (λ=0.0) or a shorter (λ=0.1) wiring length (Fig. 3B, left). These wiring lengths were chosen to represent extremes in network connectivity, allowing us to observe the effects of short and long interconnectivity on neural dynamics. As expected, increased interconnectivity, resulting from a longer wiring length, yielded attractor regions characterized by a smaller radius and lower energy values (akin to a narrow but deep basin) (Fig. 3B, left). Furthermore, a smaller radius of attractor regions likely necessitated neural states to travel a relatively longer distance to settle at the basin of these attractor regions from their initial states. That is, to achieve a stable representation, the neural states may undergo extensive transformation (i.e., heavy representation compression) (Fig. 3B, middle). To quantify this intuition, we measured the path distances, the lengths of trajectories that neural states followed to reach the basin of attractor regions, for the networks with varying levels of interconnectivity, and confirmed this observation (one-way ANOVA *F*(2,2997) = 1137.31, *p* < .001; *ps* < .001, Tukey HSD) (Fig. 3B, right).

Through this heavy compression, only the most critical features of stimuli were preserved, facilitating the achievement of higher generality. In contrast, the network with lower interconnectivity minimally transformed the representation of a stimulus by maintaining a high-dimensional space. This minimal compression allowed the stimulus to retain its unique properties crucial for achieving higher selectivity. Further formal proof (Appendix 1) confirms this visual observation that increased interconnectivity greatly compresses the representation, reducing dimensionality and enhancing generality, while decreased interconnectivity maintains higher dimensionality, enhancing selectivity.

The energy distribution in the neural manifold indicates that our model with the wiring length from the human cortex formed attractor regions that were neither too deep nor too shallow, suggesting a balanced dynamic. Accordingly, an interesting question arises regarding the characteristics of the neural network with this type of structured interconnectivity. One characteristic is stability versus adaptability of neural states in the face of perturbations arising either intrinsically (e.g., spontaneous neuronal firing) or extrinsically (e.g., top-down modulation). Neural states are less stable when neural manifolds are marked by shallow attractor regions (i.e., high energy); thus, neural states are prone to transitioning to nearby attractor regions upon encountering perturbations, indicating a propensity towards high adaptability but low stability (Gerstner et al., 2014; Izhikevich, 2007). In such case, although these networks possess high selectivity, their performance is easily influenced by perturbations, which result in chaotic behaviors. In contrast, when neural manifolds are marked by deep attractor regions (i.e., low energy), neural networks showcase resilience to both intrinsic and extrinsic perturbations (Gerstner et al., 2014; Izhikevich, 2007). However, these networks lose their adaptability, making it difficult to dynamically adjust to different task demands, resulting in ordered (stereotypical) behaviors. Our model, featuring a biologically inspired wiring length, likely straddled the boundary between order and chaos, possibly residing in a critical state. This conjecture is supported by previous studies using the Ising model, which have revealed that brain activity under resting state is near criticality (Haimovici et al., 2013; Liu et al., 2023; Priesemann et al., 2014).

To test this conjecture, we calculated the Hamiltonian value and specific heat at each temperature, using an Ising model with the same size of the network, and identified the critical temperature corresponding to the peak of the specific heat (Fig. 3C). In parallel, we used noise as input to simulate the resting-state neural activity of the network (see Methods), and calculated its Ising Hamiltonian value in comparison with that of the Ising model’s critical state. We found that the Hamiltonian value at the network’s resting state approximately coincided with the critical state of the Ising model (Hamiltonian value: -29476; critical state: -28897) (Fig. 3C). In contrast, neural networks with either a longer (Hamiltonian value: -31433) or a shorter (Hamiltonian value: -6701) wiring length resided in either ordered or chaotic regions, respectively, suggesting that deviations from this optimal wiring length propel the network away from the critical state. Therefore, our model, with its biologically constraint wiring length, operated near criticality, inherently balancing the dual objectives of perturbation sensitivity, essential for adaptability that allows the network to respond to external changes, and noise resistance, crucial for maintaining its distinctiveness and stability.

To further investigate how the criticality of our model affects its ability to recognize faces, we examined the model’s cognitive impenetrability (Pylyshyn, 1999), a signature of cognitive modules that encapsulates them from external perturbations. Specifically, two types of top-down perturbations were examined: (1) a top-down signal of faces applied in the presence of non-face objects (false alarm condition), simulating scenarios where non-face objects are incorrectly identified as faces, and (2) a top-down signal of non-face objects applied when faces were present (miss condition), representing the failure of recognizing actual faces due to interference from non-face signals. We asked whether our model was sensitive to these top-down signals and, more importantly, whether it resumed its inherent selectivity and generality after the withdrawal of these top-down signals.

For simplicity, top-down signals were exerted directly on the neurons in the model by an additional value of 2 to the neurons in the face cluster and -2 to the rest neurons to denote the presence of faces in the false alarm condition. In the miss condition, the values of the neurons in the object cluster were increased with an additional value of 2, and those of the rest neurons, including neurons in the face cluster, decreased with an additional value of -2 to denote the presence of non-face objects. In this way, the top-down signals intentionally guided the model towards a specific state (faces or objects). We then tracked the dynamics of neural activation before, during, and after these top-down signals to observe our model’s response to perturbations. In the false alarm condition, the activation of face-responsive units was first enhanced due to the top-down face signal, even in the presence of non-face objects. However, following the withdrawal of the top-down face signal, the model’s state reverted to its original dynamics, with a significant decrease in neurons’ responses to non-face objects (Fig. 3D, left), indicating that our model was able to recover its original selectivity once the perturbation was removed. In contrast, in the miss condition, the neurons’ responses to faces were suppressed due to the top-down object signal. However, the model’s state resumed its dynamics upon signal removal (Fig. 3D, right), illustrating its ability to restore its generality after the perturbation. Therefore, the top-down signals were unable to revert the network’s dynamics permanently, suggesting that our model is cognitively impenetrable, similar to the functional modules in the human VTC (de Beeck & Baker, 2010; Perrett, 1990; Pylyshyn, 1999).

## 3. Discussion

In this study, we address the challenge of balancing selectivity and generality in object recognition through a population coding perspective, proposing that the dimensionality of the neural space is crucial for achieving this balance. By examining macaques’ IT cortex, particularly the TEO and TEa, and employing computational modeling, featuring intra-module excitatory and inter-module inhibitory connections, we establish a causal link among invariance, interconnectivity, and dimensionality. Specifically, we demonstrate that proper interconnectivity among neurons created an optimal dimensionality of the neural space, allowing for efficient energy distribution across the representational manifold embedded in the neural space to balance selectivity and generality. A key aspect of interconnectivity is the wiring length derived from biological brains, pivotal in achieving this optimal balance by ensuring efficient connectivity within the network. Additionally, this wiring length facilitates the network’s maintenance in a critical state and promotes the formation of cognitive modules. In conclusion, our study underscores the role of efficient neural wiring in balancing detailed and generalized representations, providing an example of the principle in life science that structure determines functionality.

In terms of dimensionality of the neural space, selectivity and generality on object recognition can be achieved simultaneously within a unified system. In fact, in macaques’ IT cortex, the TEO and TEa achieved both selectivity and generality, and only differed quantitively in their preference, with the TEO inclined to selectivity and the TEa to generality. Importantly, this one-system view, operating at the population level, complements rather than contradicts the traditional two-system view, operating at the neuronal level. The two-system view on object recognition emphasizes hierarchical processing of objects, where selectivity is achieved through neurons tuned for features such as specific edge orientations and spatial frequencies, whereas generality is achieved through the pooling operations that combine information from multiple neurons with similar feature preferences but different positions and scales (Grill-Spector & Malach, 2004; Hubel & Wiesel, 1962; Riesenhuber & Poggio, 1999; Rolls & Deco, 2001). Our findings further demonstrate that this hierarchical structure is facilitated by interconnectivity, allowing neurons with varying sensitivities to form complex networks where selectivity and generality are inherently balanced. On one hand, interconnectivity among neurons can sharpen their tuning curves by refining responsiveness to specific stimuli and reducing reactions to irrelevant ones through mechanisms such as lateral inhibition (Blakemore & Tobin, 1972; Pinto et al., 1996), recurrent excitation (Douglas & Martin, 2004; Freeman, 1995), and synaptic weight adjustment (Bi & Poo, 1998; Markram et al., 1997). Additionally, interconnectivity can foster nonlinear mixed selectivity by enabling neurons to integrate inputs from diverse sources and respond to combinations of features rather than individual elements (Barak et al., 2013; Langdon et al., 2023; Ma et al., 2023; Rigotti et al., 2013). Neurons with either narrower tuning curves (Cai et al., 2024; De & Chaudhuri, 2023; Kim et al., 2020; Kriegeskorte & Wei, 2021; Langdon et al., 2023) or nonlinear mixed selectivity (Fusi et al., 2016; Ma et al., 2023; Panzeri et al., 2015; Rigotti et al., 2013) increase the dimensionality of the neural space, enhancing networks’ ability to encode complex and diverse stimuli to achieve selectivity in object recognition. On the other hand, interconnectivity can also reduce the dimensionality of the neural space by clustering neurons with similar response properties through Hebbian plasticity (Feldman, 2012; Hebb, 2005), pooling and integrating the inputs to compress the input space (Kanitscheider & Fiete, 2017; Marr et al., 1991), enabling functional segregation to allow different groups of interconnected neurons to specialize in processing specific types of information (Marshel et al., 2019) and recurrent inhibition to suppress irrelevant or redundant responses (Hennequin et al., 2017; Isaacson & Scanziani, 2011). These mechanisms create efficient, abstract, and generalizable representations of objects, hereby reducing dimensionality to generalize across different instances of the same object category. In summary, interconnectivity crucially modifies neurons’ response profiles, and thus dynamically regulates activation patterns of the network and its dimensionality, ensuring the network can efficiently process complex information and achieving both selectivity and generality essential for robust object recognition.

To satisfy these multiple functionalities, interconnectivity among neurons must be structured. In fact, we demonstrate that uniform connectivity in RNNs undermines the networks’ ability to develop distinct representations and hierarchical structures necessary for fine-grained object recognition. Thus, RNNs possess only generality but lack selectivity. In our model, we adopted Hebbian and anti-Hebbian rules to form structured interconnectivity. Hebbian rule reinforces the desired patterns, while anti-Hebbian rule ensures that these patterns remain distinct and non-overlapping, resulting in distinct attractors corresponding to low-energy, stable configurations (Hebb, 2005; Hopfield, 1982). Further, the analysis on the energy-representation manifold illustrates that varying levels of interconnectivity shape the attractor regions’ geometry, with higher interconnectivity resulting in narrower, deeper attractor regions that favor generality and lower interconnectivity creating broader, shallower attractor regions that enhance selectivity. The levels of interconnectivity are determined by wiring lengths, and the optimal wiring length is inspired by the wiring length of the biological brain, resulting in attractor regions in the energy-representation manifold that best balance selectivity and generality. Interestingly, this specific wiring length places the network in a critical state that balances sensitivity to perturbations for adaptability and noise immunity for stability. With this balanced adaptability and stability, a cognitive module for faces emerges, with the key characteristic of cognitive impenetrability, allowing the network to maintain its functionality despite external influences. Therefore, with structured interconnectivity derived from Hebbian and anti-Hebbian rules equipped with biologically constraint wiring length, our model provides a robust framework for efficient information processing within cognitive modules.

In summary, our study highlights the critical role of structured interconnectivity in balancing selectivity and generality by modulating the dimensionality and geometry of the neural manifold in the high-dimensional neural space. Given that interconnectivity is shaped by learning processes through mechanisms such as Hebbian learning (Hebb, 2005), synaptic plasticity (Malenka & Bear, 2004), experience-dependent plasticity (Buonomano & Merzenich, 1998; Hubel & Wiesel, 1965), dendritic remodeling (Holtmaat & Svoboda, 2009), and neurotransmitter modulation (Katz & Miledi, 1965), our study provides a populational coding perspective on how the brain adapts its neural circuits in response to new information and experiences, potentially unveiling new frameworks for understanding neuroplasticity. For example, future studies could involve training subjects on specific tasks and tracking changes in interconnectivity and neural manifold geometry over time. By elucidating the mechanisms through which learning shapes interconnectivity, we can better comprehend how cognitive functions evolve and adapt, ultimately informing the development of more effective educational strategies and therapeutic interventions for neurological disorders.

## Acknowledgments

We deeply appreciate Dr. Pinglei Bao to share the macaques’ data. This work was supported by Beijing Municipal Science & Technology Commission, Administrative Commission of Zhongguancun Science Park (Z221100002722012), and Double First-Class Initiative Funds for Discipline Construction.

## Author Contributions

Y.Z. and J.L. conceived the study. Y.Z. built computational models and performed model analyses with JR.L.’s assistance. Y.Z. analyzed macaques’ data. Y.Z. and J.L. interpreted the data. Y.Z., JR.L., and J.L. wrote the manuscript.

## 4. Materials and Methods

### 4.1 Stimuli

#### 4.1.1 Stimuli for single-unit recordings

The stimuli used for single-unit recordings in macaques’ IT cortex included 51 exemplars from six different categories: animals, vehicles, faces, vegetables, houses, and objects. Each stimulus was presented in 24 different views to capture a range of visual perspectives. The face stimuli were generated by FaceGen (https://www.facegen.com) with randomly selected parameters to ensure variability. Stimuli for other categories were sourced from a 3D model repository (https://www.3d66.com). The 24 views for each exemplar were created using 3DMAX software. For additional details on the stimulus design, see Bao et al. (Bao et al., 2020).

#### 4.1.2 Stimuli for the model

To examine face selectivity, we primarily used stimuli from the ImageNet1000(mini) dataset (Deng et al., 2009), which contains 1,000 categories with approximately 40 exemplars per category. Specifically, we selected animal faces from the cat and dog categories. Lemons were included because they mimic the external feature of faces (i.e., roundness), while vehicles, specifically ambulances and airplanes, were chosen due to their face-like configurations. To ensure diversity among non-face object categories, we randomly selected four categories: two from outdoor scenes (stores and backpacks) and two from indoor scenes (pitchers and speakers). Face stimuli were selected from the LFW Face Database (Huang et al., 2008), as the face category was not included in the ImageNet1000(mini) dataset. In total, ten categories were used for the face selectivity test (for stimulus examples, see Fig. S5).

To examine face generality, we included faces from different species (cat, dog, and tiger), randomly selected from the ImageNet1000(mini) dataset, with 10 exemplars per category. Additionally, we utilized FaceGen to generate 15 human faces from the frontal view (0°) and their corresponding profile (90°), cheek (faces rotating 135°), and back (180°) views. Importantly, the human faces used in the face generality test were different from those used in the face selectivity test, as well as from the natural stimuli used to train AlexNet, to rigorously assess the model’s generalizability. Three types of tools (vacuum cleaners, electric drills, and knives) from the HCP dataset (Van Essen et al., 2013) were also included as baseline objects. In total, ten categories (3 animal faces, 4 views of human faces, and 3 tools) were used in this test (for stimulus examples, see Fig. S5).

### 4.2 Analyses on single-unit data

#### 4.2.1 Neuron selection

Single-unit data were acquired from the IT cortex of two head-fixed male macaques as they passively viewed the stimuli. The stimuli were presented on a CRT monitor in a random order, with a screen view angle of 27.7° × 36.9° and a stimulus view angle of 5.7°. The fixation point had a diameter of 0.2°. The position of the macaques’ eyes was monitored using an infrared eye-tracking system (ISCAN) to ensure accurate fixation during stimulus presentation. Electrical signals were recorded from a total of 483 neurons in the IT cortex, with 106, 175, and 201 neurons from the TEO, TEp and TEa, respectively. Of those, 10, 190, 67, and 216 neurons were located in face-, body-, stubby-, and spiky-responsive patches, respectively. For additional details on the recordings and the characteristics of these neurons, see Bao et al. (Bao et al., 2020).

To select neurons for the analysis of selectivity and generality, we applied two specific criteria: (1) The average response of a neuron to a category had to exceed twice the maximum of the average response to other categories, ensuring that the neuron showed a strong preference for that category. (2) The number of neurons satisfying the first criterion had to be greater than five, which is the minimum required to support reliable populational coding. As a result of these criteria, only neurons responsive to the animal category met the inclusion requirements. The number of neurons responsive to other categories did not exceed five, and therefore, they were excluded from further analyses. For detailed information on the number of category-responsive neurons, see Table S1. A similar procedure was used to select spiky-responsive neurons. Spiky-ness was defined as a stimulus feature characterized by protrusion, with stimuli possessing this feature labelled as spiky stimuli (for details, see Bao et al. (Bao et al., 2020)). Of the total 51 exemplars, the spiky stimuli contained 11 exemplars, while the non-spiky (stubby) stimuli contained 9 exemplars. Each exemplar was presented in 24 different views. A neuron was defined as spiky-responsive if its response magnitude to spiky stimuli was at least twice as large as its response to non-spiky stimuli. In the replication analysis conducted on each macaque, spiky-responsive neurons were extracted from each macaque separately, following the same procedure.

#### 4.2.2 Assessment of selectivity and generality

To quantify the selectivity of animal-responsive neurons, the selectivity score was defined as the ratio of the average response magnitude of the animal-responsive neurons to animal stimuli compared to the maximum average response magnitude to non-animal stimuli. A higher selectivity score indicates greater selectivity for animal stimuli. To ensure the robustness of this measure, we randomly selected exemplars from each category 20 times, calculating a selectivity score for each iteration. By averaging these 20 selectivity scores, we obtained a final selectivity score for each animal-responsive neuron.

To quantify the generality of animal-responsive neurons, we calculated two types of generality scores. To measure generality across different animal exemplars (e.g., dogs, cats, birds, ducks, horses, and fish), the generality score was defined as the ratio of the minimum to maximum response magnitude of an animal-responsive neuron in response to all exemplars. A higher score indicates that the neuron responds consistently across different animal exemplars, suggesting greater generality. To measure generality across different views of the same exemplar, the generality score was defined as the ratio of the minimum to maximum response magnitude across different views of the same exemplar. A higher score indicates greater view invariance, meaning the neuron responds consistently to different views of the same exemplar. The calculation for both scores was repeated 20 times to obtain a reliable generality score for each animal-responsive neuron. Additionally, we used standard deviations to measure generality, as the smaller the standard deviation of neural responses to different exemplars of the same category (or different views of the same exemplar), the higher the generality.

The selectivity and generality of the spiky-responsive neuron were assessed using the same procedure.

#### 4.2.3 Effective dimensions and radii of point clouds

Chung and colleagues (2018) (Chung et al., 2018) developed a theory of linear divisibility of manifolds, which defines the effective dimension and radius of a single manifold using anchors at the edges of the manifold. Specifically, the effective dimension is defined as the spread of these anchor points along the different axes of the manifold, while the radius is the total variance of the anchor points normalized by the mean distance from the center of the manifold. To apply Chung et al.’s method to our data, we first randomly selected 100 neurons in each of the three regions (TEO, TEp, and TEa). We then used the Louvain algorithm, a method for detecting communities by optimizing a quality function known as clustering (Fortunato & Hric, 2016), to segment all neural representations into several representational manifolds. The Louvain algorithm works by iteratively moving nodes between communities to maximize clustering, resulting in cohesive manifold groups. Finally, we applied Chung et al.’s method to calculate the effective dimension and radius of each manifold. This procedure was repeated 20 times to obtain reliable values for the TEO, TEp and TEa.

For the analyses performed on each macaque, the same procedure was followed, except that only 30 neurons were randomly selected for each region due to the smaller number of neurons available. The calculation was repeated 30 times to ensure robustness in the results for each macaque.

### 4.3 Brain-inspired computational model

The computational model consists of two main components. (1) Image encoder. The first component is an image encoder, implemented using a deep convolution neural network (AlexNet), which transforms images into vector representations (i.e., object space) to serve as input for the second component of the model. (2) Simulation of human visual temporal cortex. The second component simulates the human VTC using a stochastic Hopfield-like neural network with lateral connections. Critically, the wiring length of the lateral connections is derived from the estimation of the human brain. Together, these two components enable the computational model to process images in a manner that reflects the structure and function of the human visual system.

#### 4.3.1 Recurrent Stochastic Hopfield Neural Network

In our previous study, we demonstrated that a hybrid model consisting of a pretrained AlexNet and a self-organizing map (SOM) could preserve the topological organization of fine-scale functional regions in the human VTC (Zhang et al., 2024). However, neurons in the human VTC are characterized by extensive lateral connections, particularly among neighboring neurons. These lateral connections, which can be either excitatory or inhibitory, contribute to the dynamical properties of the network, enabling complex processing and integration of visual information. Building on our previous work, in this study, we implemented lateral connections in the SOM to better simulate the functionality of the human VTC, with a particular focus on the fusiform face area (FFA), a region specialized for face processing. Importantly, by incorporating these lateral connections, our model enables a direct exploration of the causal link between structured interconnectivity, the dimensionality of neural space, and the balance between selectivity and generality in object recognition.

We incorporated excitatory and inhibitory connections between neurons in the SOM, and the weights of these connections were trained using 4 memory patterns. These memory patterns represent the activation patterns corresponding to faces, daily objects, places and bodies in our original hybrid model. In accordance with classical Hopfield neural networks, each of these patterns was binarized, assigning a value of +1 to neurons in the functionally activated areas and -1 to neurons that were not activated. This binarization process ensures that the network can store and retrieve these patterns, reflecting the distinct functional areas observed in the human brain. We use 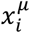 to denote the activation state of the *i*-th neuron at the *μ-*th pattern:

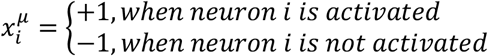

These four memory patterns serve as functional areas within the SOM, guiding the organization of neurons in the network. Lateral connections between neurons were established based on Hebbian and anti-Hebbian learning rules, which allow the network to strengthen or weaken connections depending on the correlated activity of the neurons. These rules transformed the original SOM into a recurrent neural network, enabling dynamic signal transmission across the network. Specifically, the weights between neuron *i* and *j* are calculated as a weighted sum:

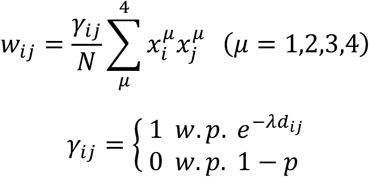

whereas the factor *γ*_*ij*_ determines whether a connection exist between neuron *i* and *j*, with *γ*_*ij*_ equal to 1 with probability 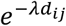 or 0 otherwise. Here, *d*_*ij*_ represents the Euclidean distance between neuron *i* and *j*. As the distance *d*_*ij*_ increases, the probability of a connection decrease, making *γ*_*ij*_ and the corresponding weight *w*_*ij*_ more likely to be 0. The parameter *λ* is a constant that determines the range of lateral connections, controlling how rapidly the probability of a connection decreases with distance and thus effectively limiting the influence of neurons on each other based on their spatial separation within the network. The value of *λ* was derived from estimation of the human brain (i.e., a biologically constraint parameter) (Theodoni et al., 2022).

In our model, we set the parameter *λ* to represent the wiring length of lateral connections in the human temporal lobe, and kept it constant throughout the model training. Though the exact value of *λ* in the human temporal lobe has not been directly measured, a recent study by Theodoni et al. (2022) estimated its value to be approximate 0.1, based on comparative studies of regions homologous to the human temporal lobe across different species. To fine-tune the parameter *λ* for our model, we aligned the human VTC with the lattice of the model. Specifically, we first calculated the half-height width of the function 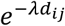 as 6.93 mm when *λ* is 0.1. Next, we computed a geodesic distance map (GDM) with a geodesic distance of 6.93 mm around each vertex of the human VTC. For both the left and right human VTC, we mapped the GDM of each vertex onto the lattice using a deformation field, and then calculated the average number of neurons in the lattice that the GDM of one vertex occupied. As a result, when the parameter *λ* in the model was set to 0.023, the number of neurons occupied by the half height and width of 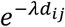 in the model approximately matched the average number of neurons occupied by the GDM of one vertex in the human VTC. Therefore, the parameter *λ* in our model was set to 0.023, closely approximating the wiring length of lateral connections in the human temporal lobe.

#### 4.3.2 Network dynamics

Dynamics are also implemented in our model to update the neural states of neurons, simulating the dynamic behavior of biological neural systems. In accordance with classical stochastic Hopfield neural networks, the state of each neuron, which can be either +1 (activation) or -1 (deactivation), is influenced by neurons to which it is connected during each iteration. That is, at each iteration, neuron *i* is randomly selected from neurons in the lattice. This neuron then updates its state *s*_*i*_ by integrating its current activation state with an input signal *b*_*i*_:

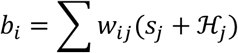

where *b*_*i*_ is a weighted sum of the state *s*_*i*_ of neuron *j* and an external input ℋ_*j*_, which represents a signal from outside the VTC that influences neuron *j*. The state *s*_*i*_ of this neuron becomes +1 (activation) with a conditional probability *p*(*s*_*i*_ = +1|*b*_*i*_), given by:

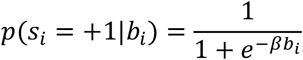

This probability is determined by a sigmoid function, where the input signal *b*_*i*_ is weighted by the parameter *β*. The magnitude of *β* controls the steepness of the sigmoid function, capturing the intrinsic randomness in the model’s dynamics. A larger *β* results in a steeper sigmoid function, meaning the system is less influenced by noise and more deterministic in its response. In our model, *β* was set to 100 to simulate a very low-level noise, ensuring that the model behaves in a highly deterministic manner. As a result, the model functions as an attractor network, capable of stabilizing around the four memory patterns corresponding to faces, bodies, places, and daily objects.

### 4.4 Invariance and corresponding neural geometry

#### 4.4.1 Selectivity and generality

To assess the selectivity and generality of the face cluster in our model, we averaged the neural states of all neurons within the cluster once the network was stabilized. Specifically, in each trial, an image from a category was processed by the model, resulting in an initial state for each neuron. The neural states were then updated over 150,000 iterations in the presence of the input image until a stable state was achieved. Our preliminary study indicated that 150,000 iterations were sufficient for the model to stabilize. To depict the developmental trajectory of the neural states, during each iteration we calculated the mean activation magnitude of the neurons within the face cluster. This procedure was applied to both selectivity and generality tests, using different image sets (see section 4.1.2). The selectivity and generality scores were calculated using the same methods as those applied to the neurophysiological data (see section 4.2.2).

For comparison, we also measured the selectivity and generality scores of a forward neural network (AlexNet, FNN_a_) and a recurrent neural network (Hopfield Neural Network, RNN_h_). Since the FNN_a_ does not inherently possess face clusters, here we decoded face responses by connecting a small neural network as a decoder to the FNN_a_ (Wen et al., 2018). This decoder consists of two layers. The first layer contains 1,000 units with a ReLU activation function, which receives a 1,000-dimensional vector output from the FNN_a_ and reduces it to a 100-dimensional vector for input to the second layer. The second layer contains 100 units and outputs 2-dimensional vector, which was used for the binarized classification of faces and non-face objects. The unit in the second layer corresponding to the classification of faces was referred to as the face unit. To train the decoder, we used natural face stimuli from the LFW face dataset (Learned-Miller et al., 2016), which contains a total of 5,749 face images, and natural object stimuli from the Caltech256 dataset (Griffin et al., 2007). To ensure that the object stimuli contained no faces and to match the number of the face stimuli, we used an in-house face-detection toolbox based on VGG-Face (Parkhi et al., 2015) to remove images containing faces from the Caltech256 dataset, ultimately selecting 5,749 non-face object images. The decoder was trained using the stochastic gradient descent optimizer (Krizhevsky, 2014). Note that all parameters of the FNN_a_ were frozen during the decoder training. Once the decoder training was complete, we evaluated the selectivity and generality scores with the activation magnitudes of the face unit in the second layer of the decoder in response to faces from the selectivity and generality dataset (see section 4.1.2). Finally, we calculated the selectivity and generality scores using the same methods as those applied to our model.

We followed the method provided by Tang et al. (2018) to construct the RNN_h_. Specifically, recurrent connections were implemented in the last layer of AlexNet (fc3 layer) to form a Hopfield network, referred to as RNN_h_. These recurrent connections were trained using images from the HCP dataset, following the Hebbian learning rule (for detailed training methods, see Tang et al. (H. Tang et al., 2018)). Once the training was complete, we decoded the RNN_h_ using images from the selectivity and generality dataset (see section 4.1.2). To obtain face-responsive and object-responsive units, we employed a support vector machine (SVM) to perform binary classification of faces and objects (for detailed analysis methods, see H. Tang et al. (2018)). After identifying the face-responsive units, we used the same the same methods as those applied to our model to calculate the selectivity and generality scores.

#### 4.4.2 Energy-representation manifold

We used the same methods as those applied to the neurophysiological data (see section 4.2.2) to obtain the effective dimensions and radii of the representational manifold formed by 1,000 neurons randomly selected from the entire neuron population. To visualize the manifold in a 2-D surface, we enhanced the traditional UMAP method by introducing a parametric approach, which utilizes a forward neural network to parametrize the non-linear mapping between the high-dimensional data space and the two-dimensional space for visualization. This approach, referred to as parametric-UMAP, offers an advantage over traditional UMAP or other manifold learning methods by enabling the learning of non-linear mappings directly from the data. Specifically, we first generated a high-dimensional neural manifold using the model with 1,000 randomly selected natural images from the ImageNet validation set (Deng et al., 2009). We then applied the UMAP to project this manifold onto a 2-D surface. In this process, a feedforward neural network was used to learn the non-linear mapping from the high-dimensional space to a two-dimensional space, resulting in the 2-D coordinates that represent the geometry of the high-dimensional manifold. This feedforward neural network consists of three layers, with the first, second, and third layer containing 1,000, 100, and 2 units utilizing the ReLU activation function, respectively.

Since our model is a Hopfield-like neural network, the states of the model evolve towards states with smaller energy, reflecting the model’s tendency to minimize its energy. To depict this dynamic characteristic, we calculated the energy for each state of the network from the initial state to the stable state using the following equation:

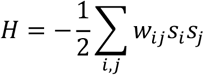

where *H* denotes the energy as a function of the network’ state, *s*_*i*_ and *s*_*j*_ are the states of neuron *i* and *j*, respectively, and *w*_*ij*_ is the connection weight from neuron *j* to *i*. Since each network state corresponds to a specific energy value, we incorporated these energy value as a third axis alongside the 2-D representational space obtained from the parametric-UMAP, creating what we refer to as the energy-representation space. This 3-D space encompasses the representational manifold and its corresponding energy values. However, traversing the entire representation space to produce a complete energy-representation manifold is impractical due to the vast number of possible states and energy values (2^40000^, as the lattice size of 200×200). To address this, we generated a proxy energy-representation manifold by performing three-dimensional interpolation, combining the energy values from 1,000 representations obtained using the parametric-UMAP method.

To calculate the path distance that neural states followed from their initial states to reach the basin of attractor regions, we recorded one representation every 10,000 iterations during a total of 150,000 iterations, resulting in 15 representations for each stimulus. We then computed the average Euclidean distance between each pair of neighboring representations within the energy-representation manifold. These distances were integrated to determine the total path distance for each stimulus. Finally, we averaged the path distances of all 1,000 stimuli used to form the manifold, providing a single measure of the path distance for the entire energy-representation manifold. This distance captures the overall trajectory of neural states as they converge towards attractor regions, reflecting the dynamics of the model.

### 4.5 Criticality and cognitive impenetrability

#### 4.5.1 Criticality estimated by Ising model

We used the Ising model to explore the criticality of our model, which is a fundamental model in statistical mechanics that describes a spin lattice system (Kardar, 2007). Each spin can be in one of two states: up or down. The model considers the interactions between neighboring spins that are first-ring adjacent, meaning they are directly connected. The Hamiltonian *H* of the Ising model represents the total energy of the system and is defined as:

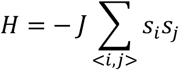

where *J* is a constant representing the interaction strength between neighboring spins (typically set to 1), *s*_*i*_ and *s*_*j*_ denote the spin states of site *i* and *j*, respectively. The summation is carried out over all pairs of adjacent spins, reflecting the interaction that contribute to the system’s energy.

The Ising model was set to have the same size as our model (i.e. 200×200) to allow for a direct comparison between the two systems. To calculate the Hamiltonian values of the Ising model, we employed Monte Carlo simulations over a range of 500 isometric temperatures, spanning from 0.001 to 10 (Landau & Binder, 2021). At each temperature, the Monte Carlo simulations were run for 1,6000,000 iterations to allow the system to reach equilibrium. Following this equilibration period, an additional 1,600,000 sampling steps were performed to obtain the average Hamiltonian value. This extensive sampling ensures that the calculated Hamiltonian values are representative of the system’s equilibrium state. Finally, the Hamiltonian values were calculated for all 500 temperatures. The specific heat was determined as the partial derivative of the Hamiltonian with respect to temperature, which revealing how the energy of the system responds to changes in temperature.

To calculate the Hamiltonian value of our model, we simulated activation fluctuations during the resting state observed in the brain by introducing random noise inputs to our model. Specifically, each neuron of the model received a time-varying noise input of either -1 or +1, following the procedure described in section 4.3.2. This procedure was used to simulate the bottom-up noise that the VTC receives from other cortical areas during the resting state. This process was run 300,000 steps, with the state of the model being recorded every 100 steps, thereby resulting in 3,000 different resting states of the model. We then calculated the Hamiltonian values corresponding to each of these states and average them to obtain the overall average Hamiltonian value for the model during the simulated resting state.

#### 4.5.2 Impenetrability of cognitive modules

We introduced top-down modulations to test the cognitive impenetrability of our model. To simulate the characteristics of top-down signals (e.g., a top-down signal of faces) observed in the brain that are briefer in time and stronger in magnitude than bottom-up signals (Bar et al., 2006; A. C. Tang et al., 2007; Wykowska & Schubö, 2010), an additional value of 2 was provided to the units in a cluster (e.g., the face cluster) and the rest neurons were provided with an additional value of -2, twice as strong as the predefined state of the model, and continuously presented bottom-up signals (i.e., the presence of stimuli throughout the simulation). Specifically, the bottom-up signals were presented for 80,000 iterations, followed by the simultaneous presence of both top-down and bottom-up signals for another 80,000 iterations. Finally, only the bottom-up signals were presented for an additional 250,000 iterations until the model reached the stable state. For every 1,000 iterations, the average response magnitude of neurons in the face cluster was recorded. This data was used to depict the developmental trajectory of the model, showing how the model evolved under the influence of both top-down and bottom-up signals.

## Supplementary information

**Table S1.** Number of neurons responsive to specific object categories or features in the TEO, TEp and TEa of two macaques’ IT cortex.

|  | TEO | TEp | TEa |
| --- | --- | --- | --- |
| Animal | 21 | 5 | 20 |
| Vehicle | 1 | 1 | 1 |
| Face | 0 | 2 | 2 |
| Vegetable | 0 | 1 | 2 |
| House | 0 | 4 | 3 |
| Tool | 0 | 0 | 2 |
| Spiky feature | 58 | 60 | 85 |

**Fig. S1.**
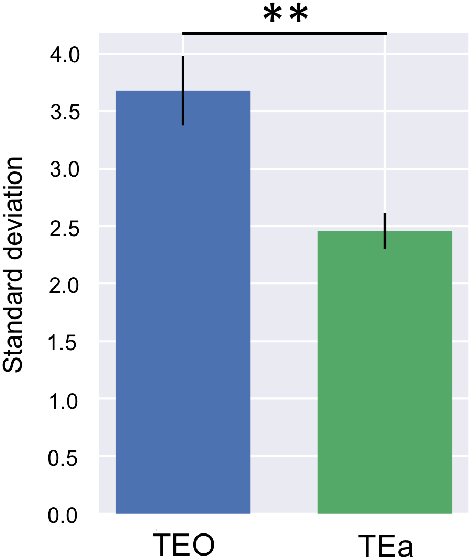
Standard deviations of neuronal responses to different animal exemplars in the TEO and TEa. The standard deviation of the responses in the TEO was significantly larger than that in the TEa (*t*(38) = 3.50, *p* < .01), indicating greater variability in the TEO’s response. **: *p* < 0.01.

**Fig. S2.**
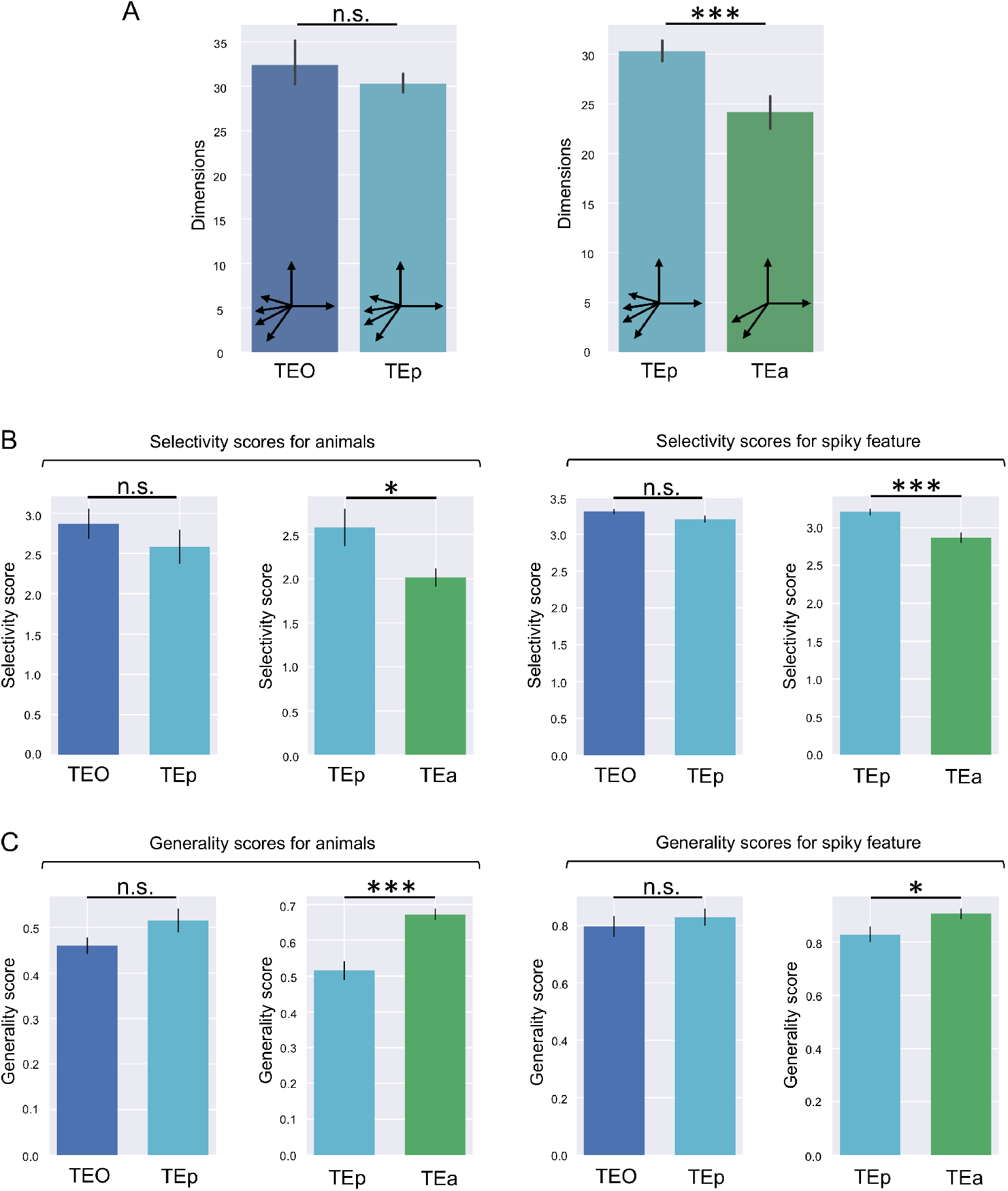
Dimensionality and corresponding invariance of the TEp compared to the TEO and TEa. Note that neurons in the TEp have a similar number of dendritic spines as those in the TEO, suggesting a similar level of interconnectivity. (A) Dimensionality. The effective dimension of the TEp was not significant different from that of the TEO (*t*(38) = -1.47, *p* = .15, top), but significantly higher than that of the TEa (*t*(38) = 6.03, *p* < .001, bottom). (B) Selectivity. The selectivity scores in the TEp for animals (versus the rest object categories, left in left panel) and for the spiky feature (versus the round feature, left in right panel) were comparable to those in the TEO (animal: *t*(38) = 0.99, *p* = .32; spiky: *t*(38) = 1.78, *p* = .08). While the selectivity scores in the TEp for animals (right in left panel) and for spiky feature (right in right panel) were significantly higher than those in the TEa (animal: *t*(38) = 2.35, *p* < .05; spiky: *t*(38) = 3.92, *p* < .001) (C) Generality. The generality scores in the TEp for animals (left in left panel) and for the spiky feature (left in right panel) were comparable to those in the TEO (animal: *t*(38) = -1.71, *p* = .10; spiky: *t*(38) = -0.75, *p* = .46). Conversely, the generality scores in the TEp for animal exemplars (right in left panel) and for spiky stimuli (right in right panel) were significantly lower than those in the TEa (animal: *t*(38) = -5.06, *p* < .001; spiky: *t*(38) = -2.28, *p* < .05). ***: *p* < 0.001; *: *p* < 0.05.

**Fig. S3.**
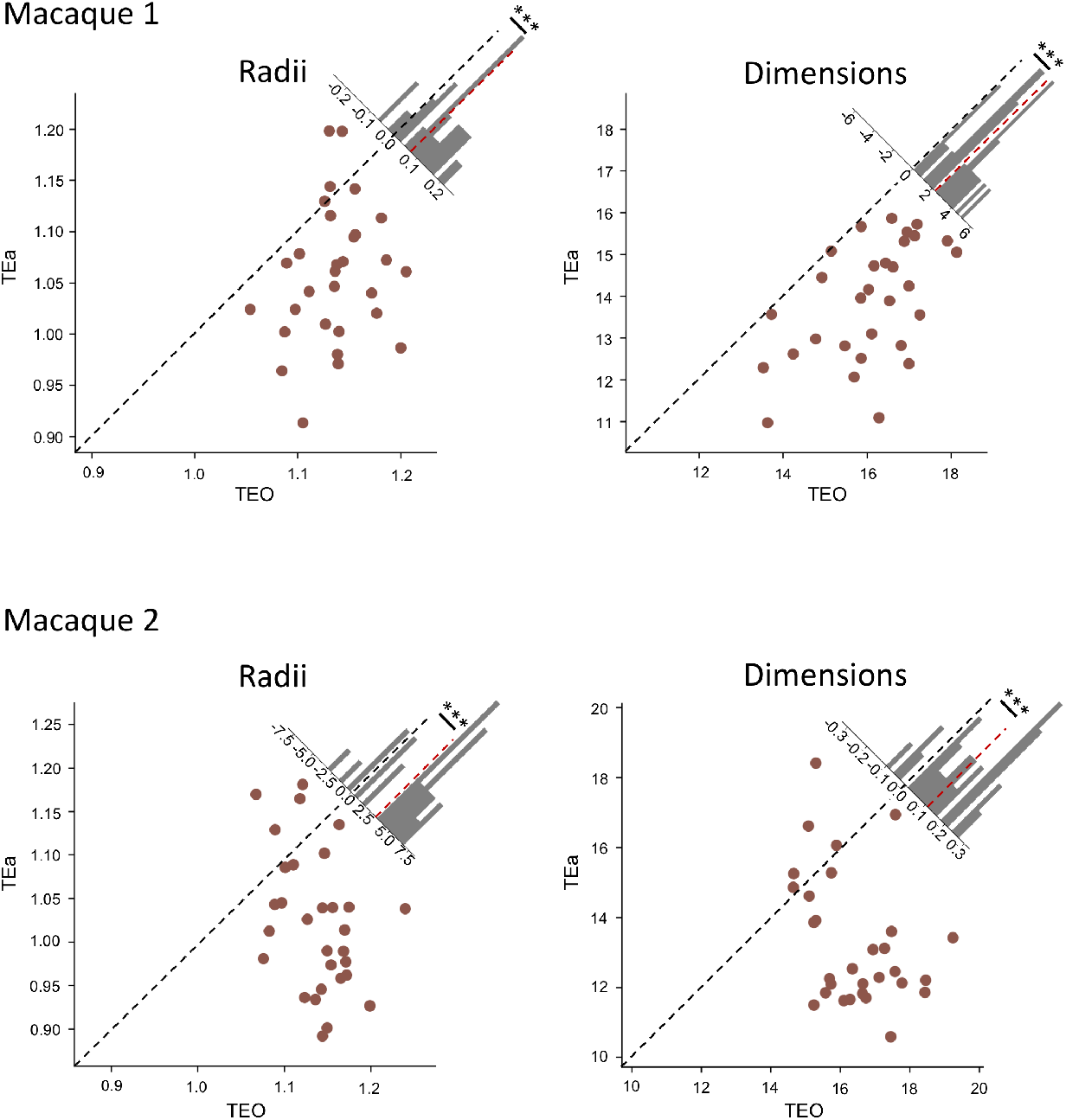
Scatterplot of TEa and TEO in two macaques, analyzed separately, with each point representing the radii (left) or dimensions (right). Points below the diagonal indicate that the dimension (or radius) of the TEO was greater than that of the TEa. The dimensions in the TEa was significantly lower than that by the TEO (macaque 1: *t*(58) = 6.41, *p* < .001; macaque 2: *t*(58) = 7.59, *p* < .001), as well as radii (macaque 1: *t*(58) = 5.64, *p* < .001; macaque 2: *t*(58) = 7.03, *p* < .001). The histogram in the upper right shows the distribution of the difference between the TEO and TEa. ***: *p* < 0.001.

**Fig. S4.**
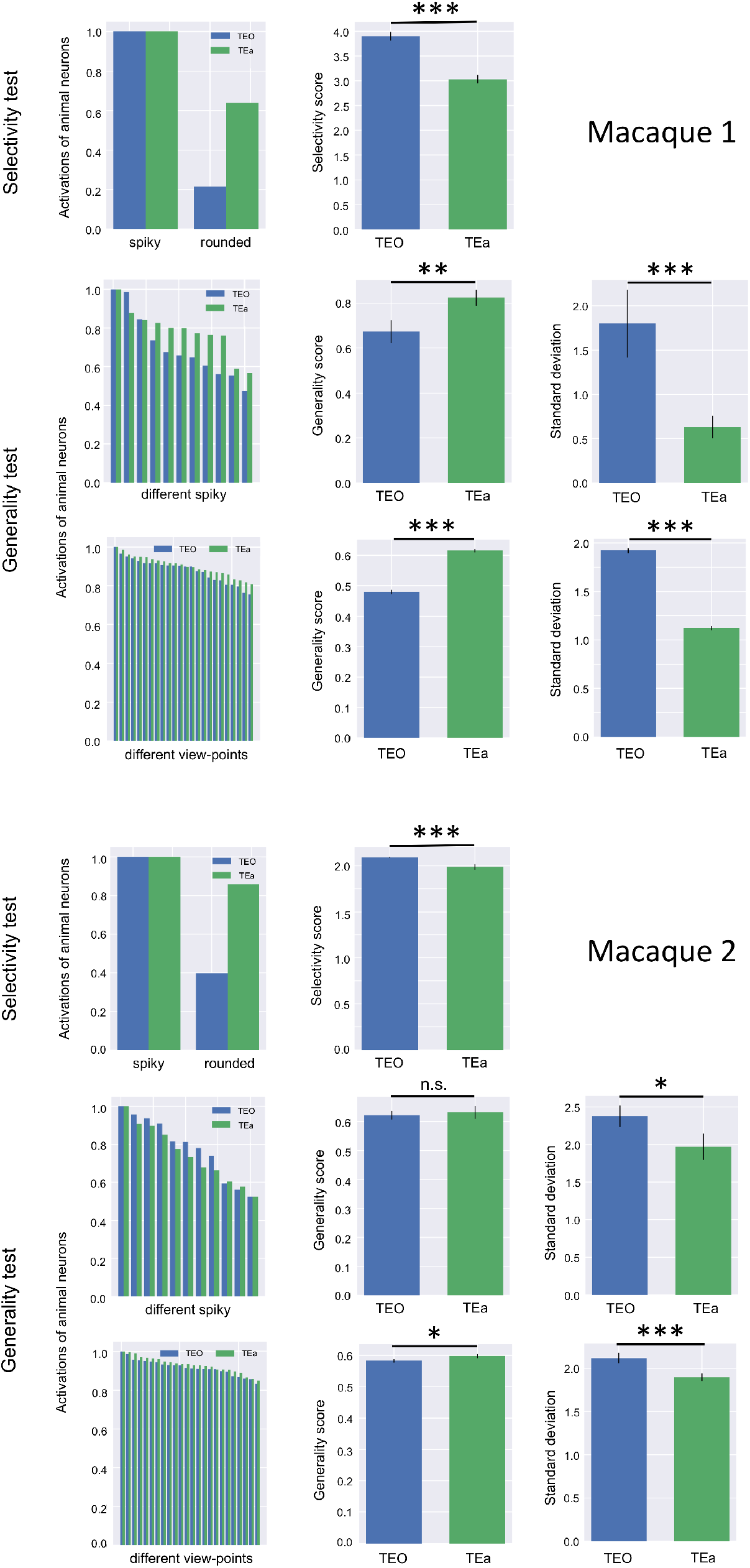
Selectivity and generality in two macaques, analyzed separately. To demonstrate the consistency of the main results across the two macaques, we conducted separate tests on the selectivity and generality of spiky-responsive neurons in each monkey. Spiky features were chosen because they are abstract and activated a sufficient number of responsive neurons necessary for the tests (Macaque1, TEO: 34, TEa: 49; Macaque2, TEO: 24, TEa: 36). In each macaque, the top row represents the selectivity test. The left panel shows the mean activation of spiky-responsive neurons in the TEO and TEa in response to spiky and non-spiky (i.e., rounded) stimuli. The right panel shows that the selectivity score of the TEO was significantly higher than that of the TEa (Macaque1: *t*(38) = 16.71, *p* < .001; Macaque2: *t*(38) = 5.05, *p* < .001). The middle row shows the generality test. The left panel presents sorted mean activations of animal responsive neurons in the TEO and TEa. The middle and right panels display the generality scores and standard deviations in the TEO and TEa. Statistical analysis shows that the TEO had lower generality than the TEa (Macaque1: generality score: *t*(38) = -3.18, *p* < .01; standard deviation: *t*(38) = 5.80, *p* < .001; Macaque2: generality score: *t*(38) = -0.83, *p* = 0.41; standard deviation: *t*(38) = 2.11, *p* < .05). The bottom row replicates the middle row with the test on generality across views of the same exemplar. Statistical analysis shows that the TEO has significantly lower generality across views than the TEa (Macaque1: generality score: *t*(38) = -20.07, *p* < .001; standard deviation: *t*(38) = 27.20, *p* < .001; Macaque2: generality score: *t*(38) = -2.24, *p* < .05; standard deviation: *t*(38) = -3.58, *p* < .001). ***: *p* < 0.001; **: *p* < 0.01; *: *p* < 0.05.

**Fig. S5.**
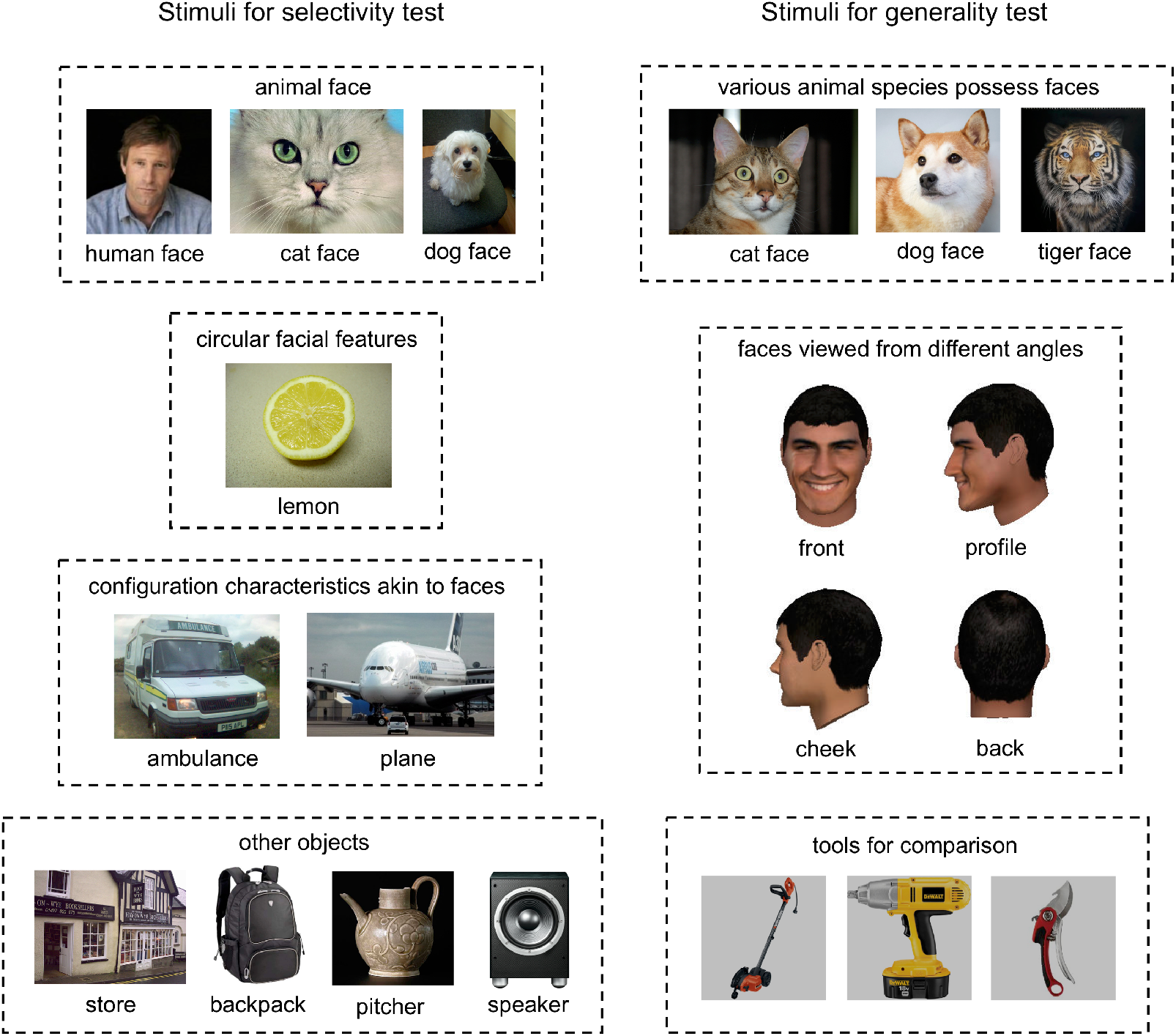
Stimulus exemplars for selectivity and generality tests.

**Fig. S6.**
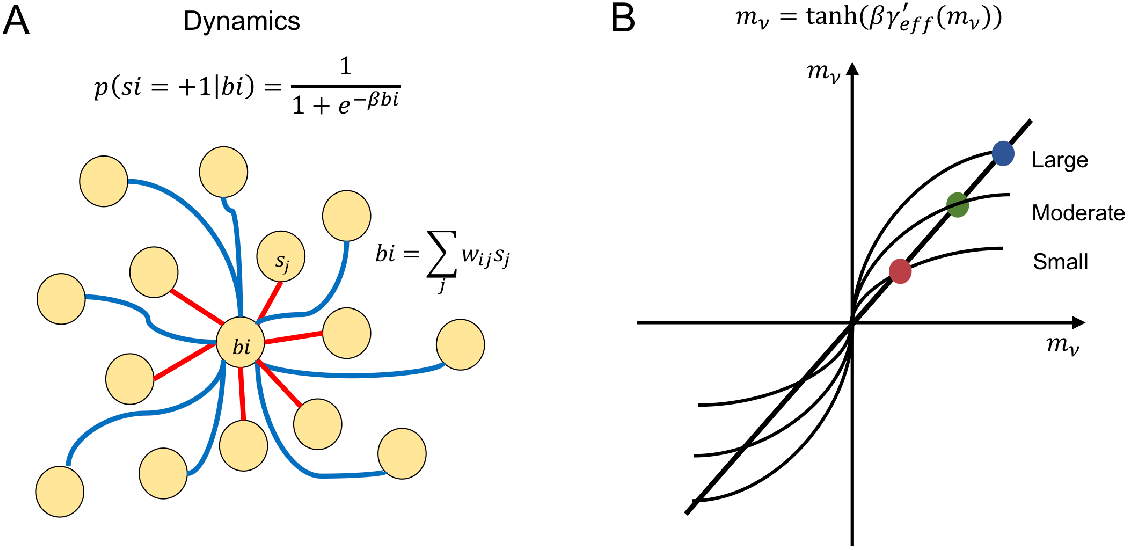
The dynamics of the model and steady states in the mean-field equation. (A) The state of unit *i* becomes ‘+1’ with a conditional probability 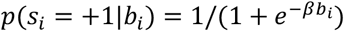. This probability function is sigmoid shaped, representing the likelihood that unit *i* will be activated under condition *b*_*i*_. The magnitude of *β* captures the intrinsic noise in the dynamics of the system. As *β* increases, indicating lower noise levels, the sigmoid function becomes steeper. For our model, we set *β* to 100, representing a very low-level of noise, ensuring that the dynamics are driven primarily by the input conditions rather than random fluctuations. This dynamical procedure is then repeated, leading the representation pattern of model to changes dynamically over time. (B) In our model, *β* is fixed, while the magnitude of 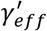, representing the interconnectivity, determines the steepness of the curve associated with the model’s steady state. As interconnectivity increases, the value of *m*_*µ*_ in the steady state also increases. A larger *m*_*µ*_ indicates a greater similarity to a specific pattern, leading to fewer possible states the network can occupy. For example, if *m*_*face*_ = 1, only one network state exists. This corresponds to a narrower attractor region, where the system is more constraint in its possible configuration, reflecting a more stable and deterministic behavior.

## Appendix: formal proof on representation compression

We utilized the mean-field technique to understand mathematical principles governing how interconnectivity modifies the depth and radius of attractor regions in the energy-representation manifold. In our model, we used the classical order parameter in a Hopfield network to quantify the degree of overlap between the network’s current state and a stored memory pattern:

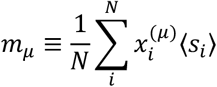

where 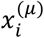 represents the state of a stored memory pattern *µ* on unit *i*,⟨*s*_*i*_⟩ is the average state of unit *i*,and *N* is the total number of units. The network’s energy can be expressed as: 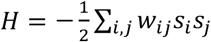, where the connection weights are given by: 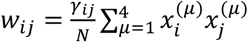. Substituting these weights into the energy equation gives:

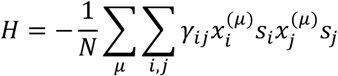

For simplicity, we replace *γ*_*ij*_ with a unified effective *γ*_*eff*_, which is proportional to the wiring length, leading to a quadratic relationship between the energy and the order parameter:

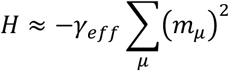

This indicates a clear quadratic relationship between the energy and the order parameter, particularly when the state is near an attractor mode (such as *s* ≈ *m*_*face*_), the energy is directly proportional to the square of the order parameter for that mode (*H* ∝ *m*^*face*^). Due to the correlation between energy and the order parameter, the order parameter can be used as a proxy for the system’s energy. Next, we investigate the relationship between the order parameter and interconnectivity.

Using the properties of two-state networks, the expression for ⟨*s*_*i*_⟩ is given by:

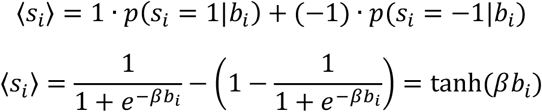

Under the mean-field approximation, *b*_*i*_ ≈ ⟨*b*_*i*_⟩, and substituting the expression for *w*_*ij*_, we obtain:

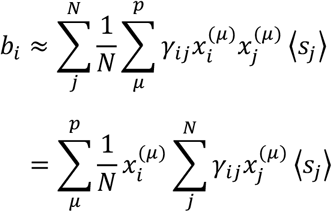

To simplify the equation, we similarly define an effective 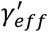 in place of each *γ*_*ij*_. The 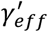 and *γ*_*eff*_ have different values but have similar properties, i.e. the larger the wiring length, the larger the 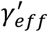. Therefore:

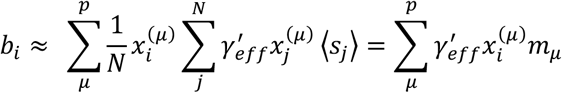

Thus, we can express the order parameter *m*_*µ*_ as:

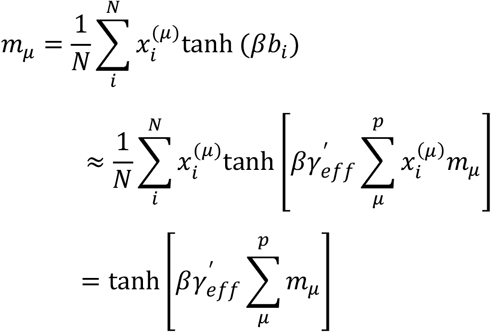

In summary, we obtain:

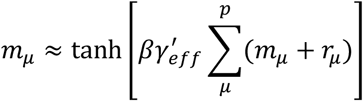

In this equation, *r*_*µ*_ corresponds to the resemblance between the bottom-up signal and the *µ*-th pattern. When the steady state of the activation pattern is close to the *v*th memory pattern, the equation simplifies to 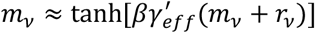. The properties of this equation, particularly the steady state of the network, can be understood by examining the intersections of these function curves (Fig. S6 B). In the model, *β* is fixed, and the magnitude of 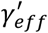, which represents the interconnectivity, determines the steepness of the curve. As the wiring length increases, the value of *m*_*µ*_ at the steady state also increases. A larger value of *m*_*µ*_ indicates a stronger similarity to the *µ*-th pattern, leading to fewer states. For example, if *m*_*face*_ = 1, only one network state exists. In this case, the attractor region is narrow. Conversely, a smaller value of *m*_*µ*_ implies more available stable states, thus corresponding to a broader attractor region.

In summary, our formal proof shows that interconnectivity plays a critical role in shaping the energy landscape of the neural network. Higher interconnectivity creates more pronounced and fewer attractor states, leading to more generalized representations. Lower interconnectivity allows for a greater diversity of stable states, supporting more selective and detailed representations. This balance between selectivity and generality is crucial for efficient neural coding in cognitive tasks such as object recognition.

